# Mechanisms involved in the production of differently colored feathers in the structurally-colored Swallow Tanager (*Tersina viridis*; Aves: Thraupidae)

**DOI:** 10.1101/2020.07.28.225862

**Authors:** Tomás Bazzano, Mendicino Lucas, Marina E. Inchaussandague, Diana C. Skigin, Natalia C. García, Pablo L. Tubaro, Ana S. Barreira

## Abstract

Non-iridescent, structurally-based coloration in birds originates from the feather’s internal nanostructure (the keratin spongy matrix), but the presence of melanin and the characteristics of the barb’s cortex can affect the resulting color. Here we investigate how this nanostructure is regulated and combined with other elements in differently-colored plumage patches. To do so, we investigated the association between light reflectance and the morphology of feathers from the back and belly plumage patches of male Swallow Tanagers (*Tersina viridis*), which look greenish-blue and white, respectively. Both plumage patches have a reflectance peak around 550 nm, but the reflectance spectrum is much less saturated in the belly. The barbs of both types of feathers have similar spongy matrices at their tips which produce similar reflectance spectra. However, the color of the belly feather barbs changes from light green at the tip to white closer to the rachis. These barbs lack pigments and their morphology changes considerably: the spongy matrix is reduced, being almost hollow, and has a different shape towards the rachis. Instead, we observed deposition of melanin underneath the spongy matrix in the back feathers which had a much saturated coloration that was consistent along the barbs’ length. Overall, our results suggest that the color differences between the white and greenish-blue plumage are mostly due to the differential deposition of melanin and a reduction of the spongy matrix in some parts of the belly feather barbs, and not a result of changes in the periodicity of the spongy matrix.

## INTRODUCTION

Plumage coloration has been traditionally split into two categories. On the one side, pigmentary plumage coloration is that generated mostly due to the deposition of pigments in the feathers. These pigments absorb specific ranges of wavelenghts of visible light and let the rest pass through, acting as light filters. Their superposition over the feather broadband-reflective structure allows the reflection of the non-absorbed wavelenghts (Shawkey and Hill, 2005), therefore generating the observable color effect through the substraction of specific ranges of visible light. The most common pigments found in the plumage are carotenoids (responsible for yellow, orange and red coloration; McGraw, 2006a) and melanins (those that generate black, gray, and the different shades of brown; McGraw, 2006b). On the other hand, plumage can be colored by the so called structural colors. Non-iridescent structural coloration is the result of coherent light scattering from the feathers internal nanostructure (known as the keratin spongy matrix), which reflects specific wavelengths and produces short wavelength colors (such as ultraviolet (UV), purple, blue, turquoise and green; Prum, 2006; D’Alba *et al*., 2012; D’Ambrosio *et al*., 2018).

However, when feathers are considered as a whole every color observed in the plumage is actually produced by an interaction of different elements, and structural and pigmentary effects can not be easily discriminated. For instance, iridescence (i.e., the change in color reflected with incidence and/or observation angle in relation to feather orientation) is considered a structural phenomenon because, in bird feathers, it is most frequently the result of constructive interference of light with a regular layered or crystal-like arrangement of melanin granules (Prum, 2006; Eliason *et al*., 2020), which absorption function is also important for the elaboration of the color effect (Shawkey and D’Alba, 2017). Also, melanin is considered to play a critical role in the production of structural coloration through the absorption of incoherently scattered white light (Shawkey and Hill, 2006; D’Alba *et al*., 2012). Without melanosomes basal to the spongy layer white reflectance would dilute the structurally reflected wavelengths leading to a whitish coloration (Shawkey and D’Alba, 2017). Moreover, a broadband reflective structure is necessary, in conjunction with the absorption of light by carotenoids, for the production of bright yellow plumage coloration (Shawkey and Hill, 2005). Consequently, the classification of plumage coloration as structural or pigmentary results arbitrary, and could be misleading when interpreting evolutionary color changes, as both types of elements are combined to produce the large array of colors observed among birds (Shawkey and D’Alba, 2017).

The study of evolution of plumage coloration is increasingly focused on understanding how the different feather elements are combined and genetically regulated (Fan *et al*., 2019; Funk and Taylor, 2019). Small adjustments in the regulatory processes leading to pigment deposition in growing feathers result in conspicuous plumage color differences between populations or species (Lopes al., 2016; Campagna *et al*., 2017; Toews *et al*., 2017). However, the regulatory processes leading to changes in structural coloration are still vastly unknown. The spongy matrix needs to be quasiordered to produce structural colors in the plumage (Prum, 2006). The wavelengths reflected by this structure depend mostly on the size and spacing between air spaces in the spongy matrix (Prum, 2006; Stavenga *et al*., 2011; Parnell *et al*., 2015), but pigments can mask or distort structurally-based color effects in the plumage (Shawkey and Hill, 2006; D’Alba *et al*., 2012; Tinbegen *et al*., 2013; Fan *et al*., 2019). It is likely that large part of the variation observed in structurally-based plumage coloration is actually due to variation in pigment deposition rather than to alterations of the spongy matrix nanostructure. Thus, to know which are the alterations that lead to changes on plumage coloration it is necessary to investigate the characteristics of the elements involved and their specific contribution to plumage reflectance.

The Swallow Tanager (*Tersina viridis*) is a sexually dichromatic South-American passerine species. While males are mostly greenish-blue, females are mostly colored green (Barreira *et al*., 2008). Males also show a white plumage patch in their bellies (Figure 1a) that appears light yellow in females. Juvenile males go through a transition from female-like plumage in their first year, to male adult plumage that takes about four years (Hilty, 2020). The greenish-blue plumage coloration of males is the result of light reflected by the quasi-periodic nanostructure present in their feather barbs (D’Ambrosio *et al*., 2017; 2018). Also, the greenish-blue plumage of male Swallow Tanagers changes with viewing geometry from green-turquoise, when the angle between the observation and illumination directions is small, to UV-blue when this angle increases (Barreira *et al*., 2016). This angle-dependent color appearance is also the result of the interaction of light with the spongy matrix (Skigin *et al*., 2019). The morphology and reflectance properties of the white belly plumage patch were not described before. White plumage coloration (i.e., lacking pigment deposition) is ideal to study the contribution of feathers’ nanostructure to their reflectance properties in bird species showing both, white and structurally-based plumage coloration. Our objective is to gain a better understanding about the mechanisms underlying color variation between plumage patches. We hypothesized that the differences in coloration between greenish-blue and white plumage patches of male Swallow Tanagers are mainly the result of the addition (or subtraction) of melanin granules in their feather barbs. Therefore, we expect to find a similar spongy matrix nanostructure in both feather types. We also aim to analyse the variation of other structures of the feather barbs (i.e., shape, cortex thickness, pigment presence and location, etc) in order to investigate how these contribute to plumage coloration. We performed a series of reflectance measurements for different configurations (e.g., measuring light reflectance on specific areas of the feather barbs and over the plumage) and characterized the feather’s morphology through transmission and scanning electron microscopy (TEM and SEM, respectively) to associate the characteristics of the reflected light directly to the morphology and the disposition of the elements present in the feathers. By comparing differently-colored feathers of the same organism we expect to achieve a better understanding about how feathers’ color-producing elements are modulated to generate different optical effects.

**Fig. 1.**
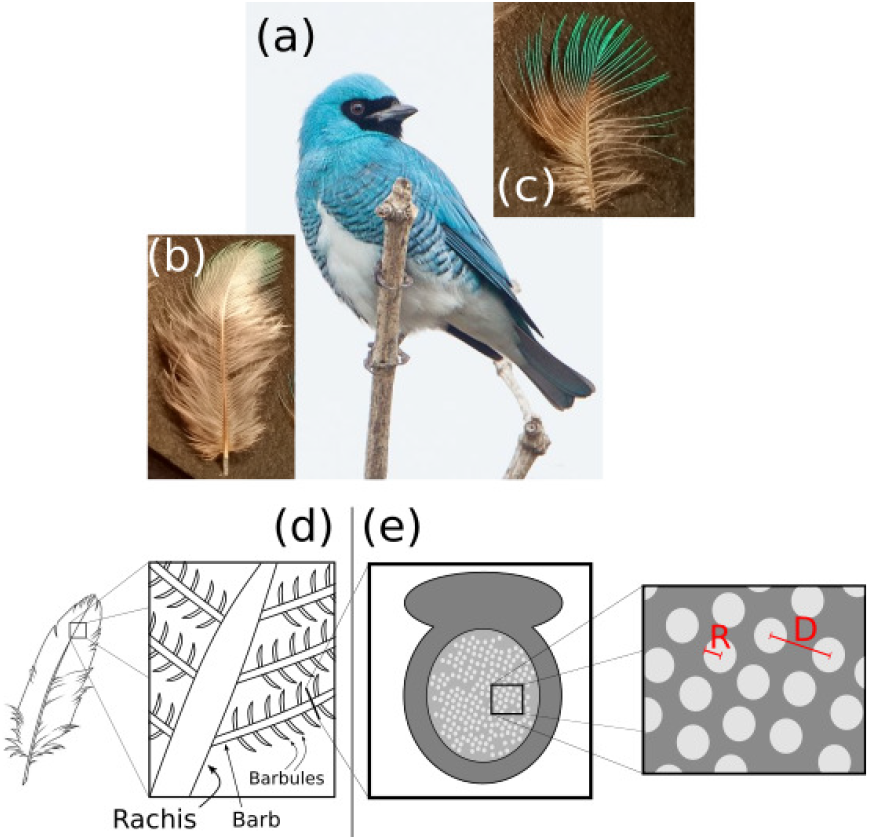
(a) Male Swallow Tanager (photo by Nortondefeis - Own work, CC BY-SA 4.0, https://commons.wikimedia.org/w/index.php?curid=49783199); (b) image of a “white” belly feather; (c) image of a greenish-blue back feather; (d) scheme illustrating the main parts of a feather; (e) diagram of the barb’s cross section, the dark gray area represents the barbs cortex and the interior area represents the spongy keratin matrix. To the right, detail of the spongy matrix where R = radius of the air vacuoles and D = distance between air vacuoles.

## MATERIALS AND METHODS

### Specimen collection

We investigated the color properties and feather morphology of male Swallow Tanagers (*Tersina viridis*) specimens deposited at the Ornithology Collection of the Museo Argentino de Ciencias Naturales “Bernardino Rivadavia.” We extracted feathers from the belly (BL, Figure 1(b)) and back (BK, Figure 1(c)) plumage patches. The feathers exhibit a colored region at their tips (Figure 1 (b) and (c)), which looks light green and occupies a small area in the BL feathers while approximately half of the BK feathers’ total area exhibits greenish-blue color. Feathers are formed by a main shaft called the rachis. Fused to the rachis are a series of barbs (where the color is produced); and the barbs themselves are also branched and form the barbules, as indicated in Figure 1(d). Within the feather barbs lays the spongy matrix of non-iridescent structurally-colored feathers (Figure 1(e)), which is formed by air vacuoles immersed in a keratin matrix. This nanostructure is surrounded by the cortex, a solid layer of keratin with variable width. The radius of the air vacuoles (R), the distance between them (D), and the thickness of the spongy matrix are determinants of the photonic response of the coloration of the BK feathers of male Swallow Tanagers (D’Ambrosio *et al*., 2017; 2018; Skigin *et al*., 2019).

### Reflectance measurements

We performed a series of reflectance measurements to describe the color properties of the feathers from the BK and BL of male Swallow Tanagers at different scales. First, we measured reflectance spectra of the BK and BL plumage patches on a Swallow Tanager museum specimen to obtain a description of the plumage coloration (as a macroscopic effect of the superposition of several feathers). Then we performed micro-spectrophotometric measurements to describe the pattern of light reflected specifically by different parts of the feathers (i.e., by individual feather barbs and directly on the spongy matrix). We describe these methods in detail next.

### Plumage reflectance

For the measurement of plumage reflectance we employed an Ocean Optics USB4000 spectrophotometer with a LS-1-LL halogen light Ocean Optics (emission range 420 to 780 nm) calibrated against a WS-1 white diffuse reflectance standard (Ocean Optics, Dunedin, Florida, USA). We measured each spectrum as the average of a number of readings between 5 and 20, with an integration time of 2000 msec, with a boxcar smoothing function of 29 points. With a bifurcated optic fiber probe, placed in a rubber holder to isolate its tip from ambient light, we illuminated the surface and collected the reflected light to determine the percentage of light reflected. We positioned the probe perpendicularly over the surface at a distance of approximately 2 mm. We measured both plumage patches 15 times each, lifting the probe and re-positioning it again in the area of interest each time, and then calculated mean reflectance spectra for each plumage patch.

### Micro-spectrophotometry of feather barbs

We measured the reflectance of single feather barbs from each plumage patch at approximately normal incidence to describe the specific contribution of a single feather barb to the overall color. We measured both, the exposed (i.e., the side that is visible) and the inner side (i.e., the side that faces the bird’s body) of BK and BL feather barbs (Figure 2(a), configurations (1) and (2)). We measured multiple times (between 5 and 22) on at least 4 different feather barbs in each feather and then calculated the mean reflectance for each side. We also measured the reflectance directly on the spongy matrix, as schematized in configuration (3) of Figure 2(a).

**Fig. 2.**
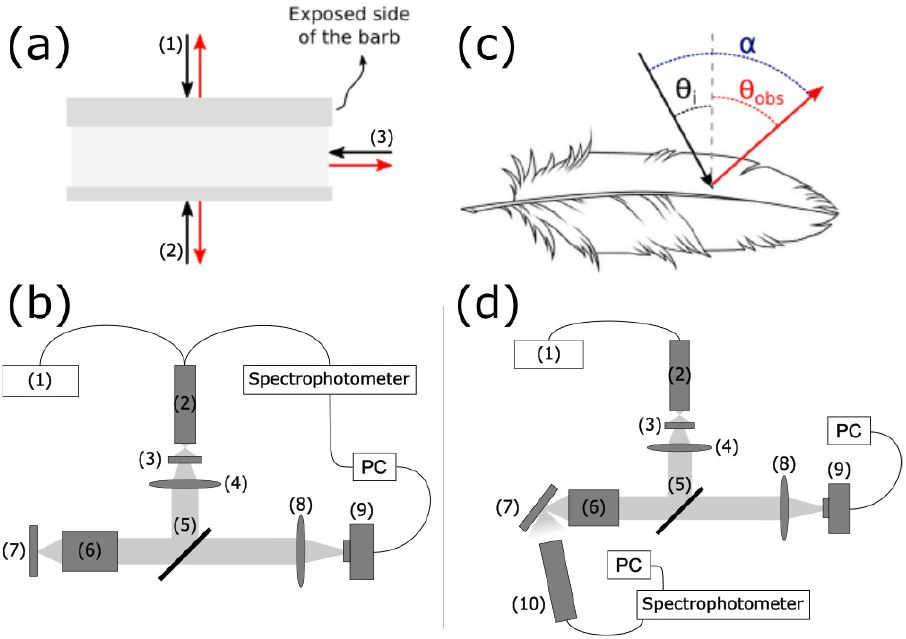
Schemes of the different types of micro-spectrophotometry reflectance measurements performed. (a) Representation of the reflectance measurements performed on the (1) exposed side of a feather barb, (2) inner side of a feather barb, and (3) spongy matrix taken from cross sections of the tips of feather barbs; (b) experimental setup used to collect reflectance data over individual feather barbs, (1) light source, (2) bifurcated optic bundle, (3) iris diaphragm, (4) convex lens, (5) beam-splitter, (6) microscope objective, (7) sample, (8) convex lens and (9) camera; (c) angle configuration used to measure the reflectance spectra with different illumination/observation geometries; (d) setup employed to measure reflectance of whole individual feathers at arbitrary incidence and observation angles, (1-9) same as for (b), and (10) fiber optic bundle.

To obtain the reflectance spectra of individual barbs under normal incidence we employed the experimental setup shown in Figure 2(b). Light is emitted by the light source through a bifurcated fiber optic bundle, that after focusing generates a spherical wave-front. Only the central part of this front passes through the iris diaphragm, to avoid chromatic aberration due to the biconvex lens that follows the path. This lens has a focal distance of *f* ~ 10 *cm* and its focal point matches the fiber optic bundle’s focal point, so that, ideally, the light beam that emerges from the lens is parallel to the optical axis. Next, light reaches the beamsplitter *Standa 081746*, which divides it into two. One of the beams passes through a microscope objective, which focuses the light on the sample. Light is reflected by the sample and passing through the objective reaches the beam-splitter back. The reflected beam returns back to the bifurcated bundle following the same path. Light that reaches the bundle is detected by the spectrometer and analyzed in the computer. On the other hand, the light transmitted by the beam-splitter passes through a secondary biconvex lens of focal distance *f* ~ 10 *cm*, and by using a camera *Infinity 1-2C*, we could obtain an image of the illuminated spot. The camera is focused on the surface of the sample, therefore, by illuminating the sample from behind, it works as a microscope, and allows to set the sample in the right position. This extra illumination was turned off during reflectance measurements. We took measurements relative to a WS-1 diffuse white standard (Ocean Optics, Dunedin, Florida, USA). We used an Ocean Optics USB4000 spectrophotometer with a halogen light Ocean Optics LS-1-LL (emission range 420 to 780 nm). Individual barbs and barbules were illuminated using a microscope objective of numerical aperture *NA* ~ 0.82 (*Olympus PLCN100×0*), resulting in a light spot of 1.27 ± 0,02 *μ*m of diameter.

### Micro-spectrophotometry with varying angle configurations

In previous studies (Barreira *et al*. 2016, Skigin *et al*. 2019) we described how the greenish-blue color of the BK plumage of male Swallow Tanagers changes from green to blue (i.e., from a longer to shorter wavelength reflectance peak) when the angle between the illumination and observation directions increases. Here, we used a micro-spectrophotometric approach to describe this effect at the individual feather. In Figure 2(c) we schematize the angle configurations used to study the reflectance of the sample for different illumination and observation angles over a feather, and in Figure 2(d) we show the setup employed for the measurements. We rotated the sample to allow different incidence angles. The light reflected by the sample was collected by a secondary fiber optic bundle, pointing at the illuminated surface so that the light reflected by the sample enters the bundle and reaches the spectrometer. In this way, the incidence angle (*θ_i_*) and the observation angle (*θ_obs_*) in Figure 2(c), could be varied. We explored 22 configurations producing 8 values of *α*. We compared reflectance spectra for different angle combinations with the same *α* since our previous results showed that the spectrum of reflected light of the plumage depends mostly on *α*, independently of the values of *θ_obs_* and *θ_i_*. In the case of non-normal illumination on samples formed by groups of a few barbs and barbules, we used an objective of *NA* ~ 0.1 (*Olympus PLCN4x*), which produces a light spot of 33 ± 3 *μ*m of diameter. For each configuration, we measured reflectance several times and calculated its mean value and the standard deviation.

### Examination of feathers structure

To identify the morphological characteristics of the feather that contribute to the color response, we examined the feather structure using electron microscopes. We obtained images of the cross sections of the barbs using a scanning electron microscope (SEM Carl Zeiss NTS-SUPRA 40) and a transmission electron microscope (TEM JEOL-100 CX II). To obtain the SEM images, the samples were previously subjected to an Au sputtering treatment of 5 - 10 nm.

We made ultra-thin cuts of barbs from the colored area of the feathers to describe the structures present through TEM images. Samples were fixed on glutaralheyde 2.5% in a phosphate buffer 0.067M (pH 7.2). The samples were then rinsed 3 times with phosphate buffer. We added osmium tetroxide 2% on destilled water for 1 hour, and made 4 washes (15 minutes each) with destilled water. Samples were then dehydrated with a gradient of ethanol/destilled water of 30 and 50% v/v (15 minutes each) and left overnight at 70% v/v. The samples were taken to ethanol 96% and then continued dehydrating with a gradient of acetone/ethanol (50% v/v, 15 minutes each) until 100% acetone was reached, where samples were left for 1 hour. Afterwards, the samples were infiltrated with a Spurr/Acetone resin at room temperature: 1:3, 1:1 v/v (1 hour each) and 3:1 v/v (overnight). The samples were then left on pure resin for a day. The resin was polimerized at 70° C overnight. Ultrathin cuts were obtained with an ultramicrotome (LKB, 2088), deposited on copper grids (300 mesh) and contrasted with uranyl acetate (1 minute) and lead citrate (5 minutes). We obtained images of these cuts using several magnifications to examine in detail the structures present. We used these images and TEM images of BK feathers we studied previously (Barreira, 2011; D’ Ambrosio *et al*., 2018).

We used the ImageJ software to measure the morphology of the barb outlined in Figure 1(e). Basically, the cross section of the barb shows an outer cortex that surrounds a keratin spongy matrix formed by quasi-spheroidal air cavities. Here we used microscopy images to measure the radius of the cavities (R) and the separation between them (D) for both BL and BK feather barbs. We estimated average values and the standard deviation of these geometrical parameters that were shown to be key features in the determination of color production in feathers (D’Ambrosio *et al*., 2017; 2018, Skigin *et al*., 2019).

## RESULTS

### Reflectance measurements

#### Plumage reflectance

Figure 3 shows the reflectance curves for greenish-blue (BK) and white (BL) plumage patches measured directly on the male Swallow Tanager museum skin at an approximately normal incidence. It can be observed that the curve for the BL patch exhibits an unsaturated reflectance peak at λ ≈ 550 nm, despite appearing white to the human eye. For larger wavelengths, the behavior is quite uniform, and for shorter wavelengths the intensity of the reflectance decreases towards the UV zone. The reflectance spectra of the BK plumage patch has a saturated reflectance peak also at λ ≈ 550 nm; but reflectance is low for larger wavelengths while it slightly increases towards the lower wavelengths range.

**Fig. 3.**
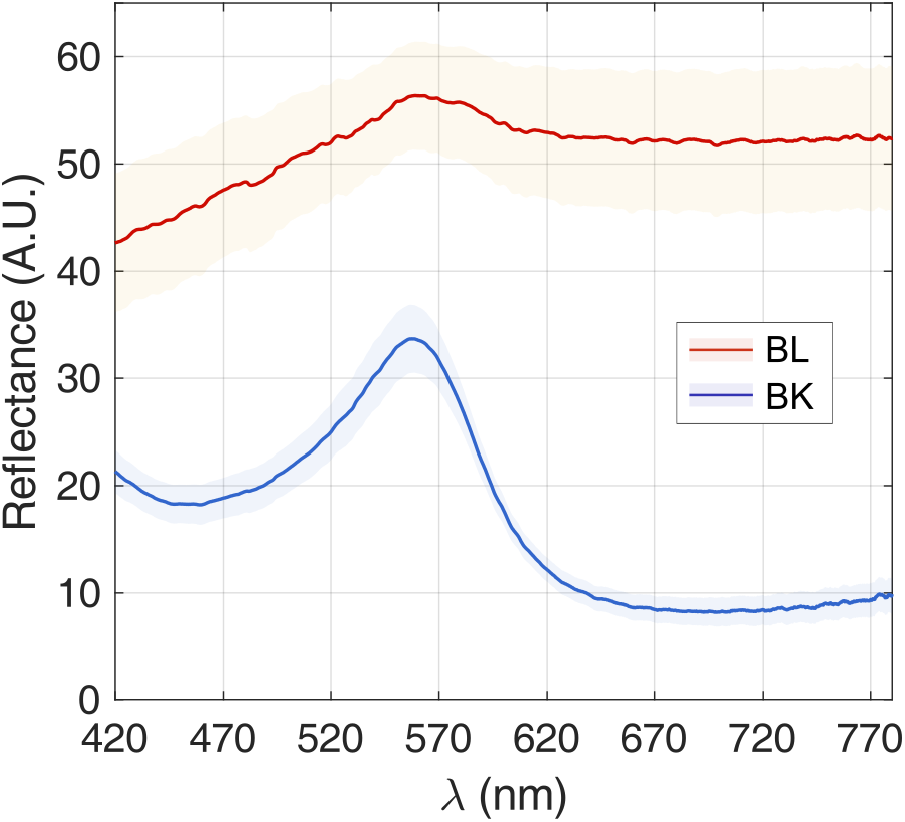
Mean reflectance spectra taken with normal incidence over the belly and back plumage patches of a museum specimen of male Swallow Tanager. Shaded areas represent standard error of each reflectance curve.

#### Micro-spectrophotometry of feather barbs

The spectra shown in Figure 3 include the collective response of several feathers, and consequently, of a set of superposed barbs and barbules, which appear with different orientations. However, a close observation of a feather under a microscope evidenced that the color is mainly generated within the feather barbs. When carefully inspecting a BL barb from the feather’s tip, we observed that it exhibits a color gradient: it is greenish near the barb’s tip and grayish white near the rachis (see Figure 1(b)).

In Figure 4 (a) and (b) we present the reflectance for individual BL and BK barbs, measured at five different positions along the barb separated ≈ 1 mm, between the rachis (A) and the tip of the barb (E), as indicated in the inset of Figure 4. These reflectance measurements were taken on the exposed side of the feather barbs. The reflectance spectra of the BL barbs showed a peak at approximately 525-530 nm for positions B-E, much more saturated than the reflectance peak obtained when measuring the complete plumage patch (Figure 3). The reflectance corresponding to position A (i.e, nearest to the feather’s rachis) showed a very uniform behavior for the entire visible range and a much lower intensity than in the previous cases. This spectrum is consistent with the white color observed in this part of the barb. Notice that the maximum reflectance intensity corresponds to measurements at position C. On the other hand, all the curves corresponding to the BK barb are more similar to each other, not only in the reflectance peak position but also in intensity, including that corresponding to the closest position to the rachis. For positions A and B, the peak is at ≈ 530 nm, and for positions C, D and E the peak is at ≈ 550 nm. We also observed that the reflectance peaks corresponding to the BL barb are wider (i.e., less saturated) than the BK ones, in agreement with the color observed at the tips of these feathers.

**Fig. 4.**
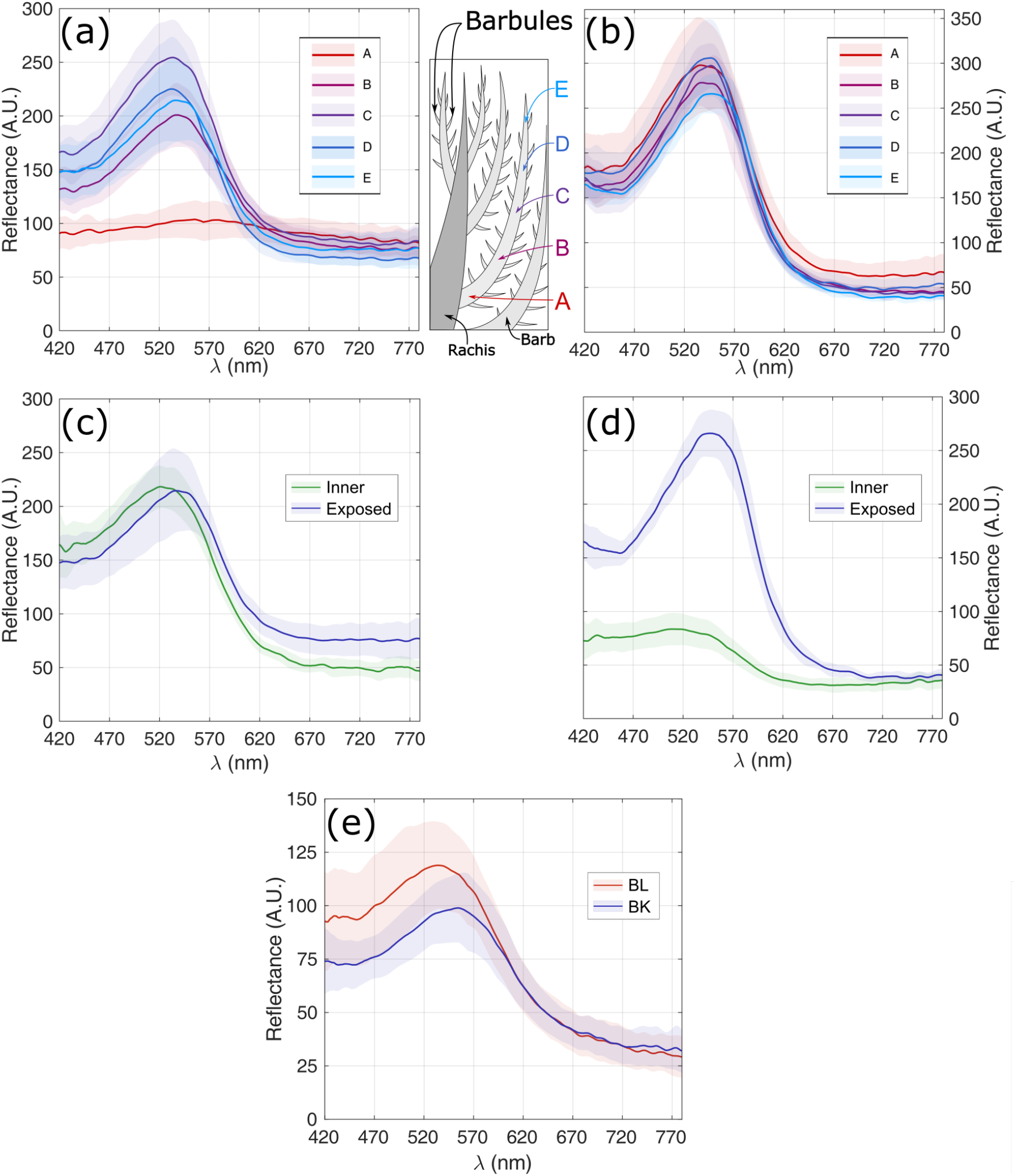
Mean reflectance spectra and standard error (shadowed area) of individual feather barbs at normal incidence. (a) Reflectance along different positions on an individual BL feather barb; (b) reflectance along different positions on an individual BK feather barb; the inset between (a) and (b) schematizes the areas of the feather barbs where reflectance measurements were taken; (c) reflectance measured on the exposed and inner side at the tip (position E) of the BL feather barbs; (d) reflectance measured on the exposed and inner side at the tip (position E) of the BK feather barbs; (e) reflectance spectra measured directly on the medullary section of the tip of BL and BK feather barbs.

Figures 4 (c) and (d) show the reflectance obtained at the tip (position E) for BL and BK feather barbs. In each case there are two curves: one corresponds to the response of the exposed side of the barb (Fig. 2(a), configuration (1)) and the other to the inner side (Fig. 2(a), configuration (2)). In the case of the BL feather, the curves (exposed and inner side) are similar; however, the peak corresponding to the inner side is slightly shifted towards the blue wavelengths. In the case of the BK feather barbs, the curves for the exposed and inner sides of the feather barbs differ substantially. The reflectance for the inner side is low and fairly uniform, with a slight intensification for wavelengths between 420 and 550 nm. Unlike the behavior of the curves for the BL feather barbs, for the BK feather barbs the measured response depends strongly on the illuminated side of the barb. This would suggest that in this case the barb presents some asymmetric characteristic among both sides that affects its light reflectance properties.

The curves shown in Figures 4 (a) to (d) include the collective response of all feather barb components, including the cortex, the spongy matrix, and pigments (if present). To investigate the individual response of the spongy matrix of the BL and BK feathers, we performed reflectance measurements at normal incidence on the transversal cut directly on the spongy matrix of each feather barb using the configuration (3) described in the scheme of Figure 2(a). Figure 4 (e) shows the reflectance spectra of the spongy matrix at the tip of BL and BK barbs. The behavior of both curves is similar. The reflectance peak for the BL feather is at λ ≈ 530 nm and for the BK feather is at λ ≈ 550 nm. Comparing the reflectance of the spongy matrix with that of the complete barb illuminated from the exposed side (Figure 4 (c) and (d)) for both the BL and the BK feathers, we observed that the spectral positions of the peaks coincide. Therefore, these results suggest that the peaks observed in the response of the whole barb (at the exposed side of the feather) are mainly determined by the interaction of light with the spongy matrix. Regarding the overall behavior of the curves, it is observed that for the BL feather there are no significant differences between the measurements performed on the spongy matrix and the complete feather barbs (except for position A), suggesting that the barb’s cortex has a non significant effect on the optical response. However, for the BK feather, the reflectance for wavelengths outside the peak decreases more abruptly in the case of the full barb measurement. This suggests that other elements present in the BK feather barbs have an influence, in addition to that of the spongy matrix, on the optical response than those of the BL feather barbs.

#### Micro-spectrophotometry with varying angle configurations

We analysed the angular behavior of the reflectance spectra for BK and BL individual feathers. We show the reflectance measurements for different angles of incidence and observation made on the colored-tip of the feathers on Figure 8 (Supp. Mat). For both, BK and BL feathers, the reflectance peak shifts towards the blue wavelengths as *α* increases (Figure 5). It can also be observed that this behavior is independent of the specific values that *θ_i_* and *θ_obs_* take, as different configurations with equal *α* values resulted in similar reflectance spectra (Figure 8, Supp. Mat). This phenomenon gives rise to a particular color response, changing from a green to a bluish hue as *α* increases for the BK feathers. A similar behavior is observed in the case of the colored-tip of the BL feathers, although the observed color in this case is much dimmer (Figure 5). Notice that for all the values of *α*, the peak wavelengths of the BK feather are slightly larger than those of the BL feather.

**Fig. 5.**
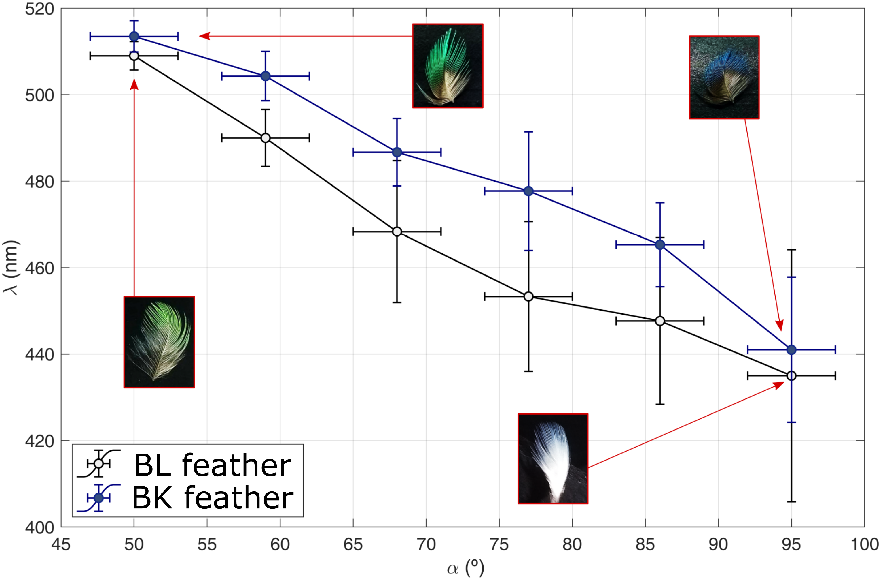
Spectral position of the reflectance peak (hue) vs. *α* for the BL feather (black line) and for the BK feather (blue line). Error bars are also included. The insets show photographs of the BL and the BK feathers for *α* ≈ 50° and ≈ 95°. For each value of *α*, the peak position was obtained by averaging the values found for the different setups (Figure 8). The error in the determination of *α* is ±3°.

### Examination of feathers structure

In Figure 6 we show SEM and TEM images of the barbs’ cross section. The shape of the cross section of both barbs is different: the barb of the BL feather has an approximately circular shape, while the barb of the BK feather exhibits a more triangular section, with a vertex at the bottom (inner side). On the other hand, the width of the cortex surrounding the spongy matrix also varies from one feather to another. The cortex thickness of the BL barb is wider in the exposed side, its value is ≈ 5 *μ*m at the tip and also near the rachis. In the BK case, the cortex thickness at the barb’s tip is more uniform around the barb, with a value of ≈ 5 *μ*m. As observed in panels (e) and (f) of Figure 6, the cross section of the BL barb varies significantly along the barb. In the mid section of the feather barb (Figure 6(e)) the external cortex thickens around the barb in relation to the spongy matrix area. Near the rachis the BL has a flatter shape and exhibits big holes with a very reduced area occupied by the spongy keratin matrix. This structure is similar to that described by Igic *et al*. (2018) for the white feather barbs of *Carduelis hornemanni*.

**Fig. 6.**
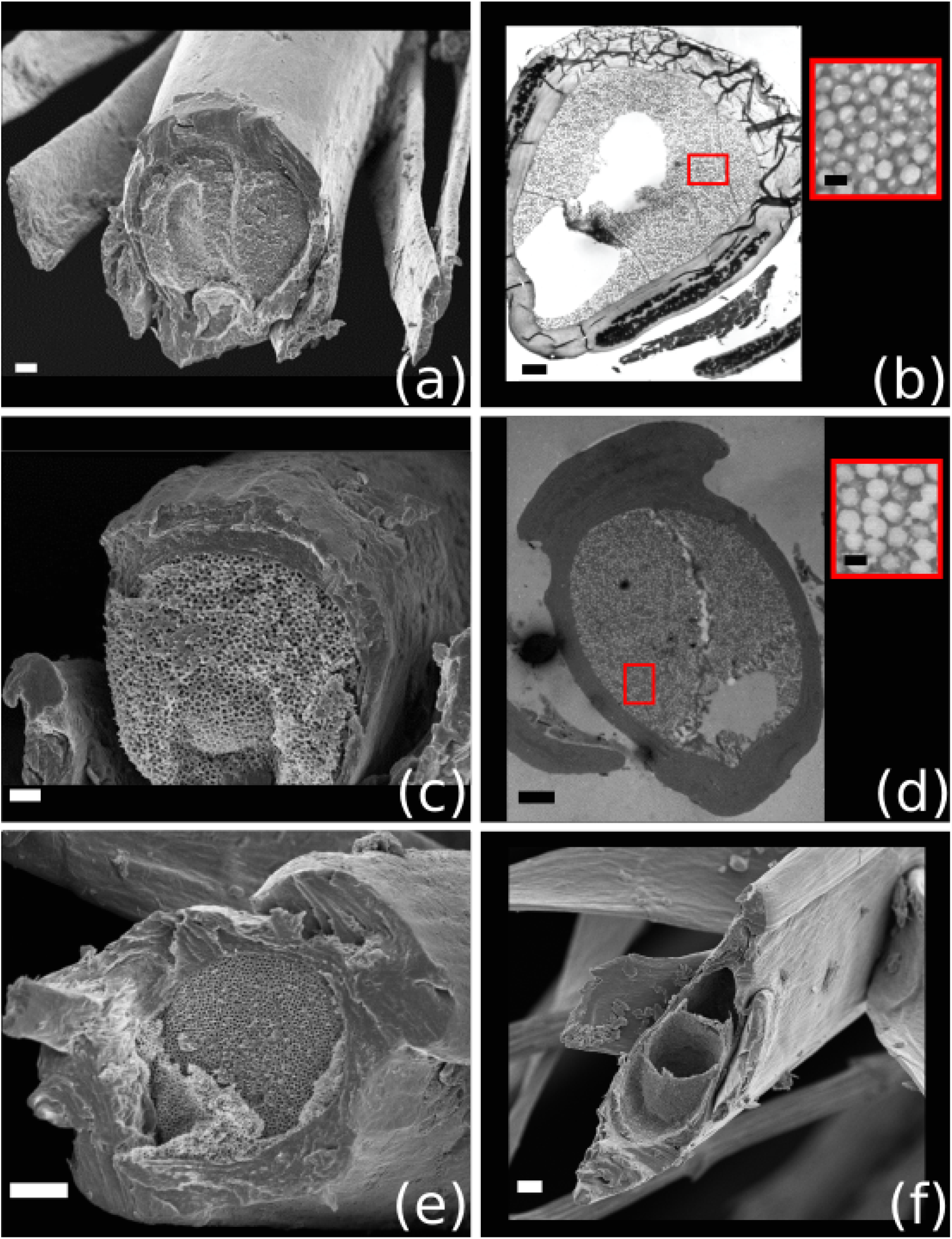
SEM and TEM images of transversal cuts of individual feather barbs. SEM (a) and TEM (b) image of the tip of a BK feather barb; SEM (c) and TEM (d) image of the colored-tip of a BL feather barb (position E); SEM image of the cross section of a BL feather barb at position C (e) and at position A (f). The insets in panels (b) and (d) show a detail of the corresponding spongy matrix. Scale bars: (a) 2 *μ*m; (b) 1.25 *μ*m, 0.2 *μ*m (inset); (c) 1 *μ*m; (d) 2 *μ*m, 0.2 *μ*m (inset); (e)-(f) 3 *μ*m. Image (b) was also published in (Barreira, 2011 and D’Ambrosio *et al*. 2018).

A fundamental difference between the BL and BK feathers is the presence of melanin granules in the lateral and inner sides of the cortex of the BK barbs and also in the barbules, in contrast to the BL feather, in which no pigments were found (Figure 6 (b) and (d), respectively. In both feather barbs, the cortex surrounds a spongy structure formed by air cavities in a keratin matrix. Previous studies on the male Swallow Tanagers greenish-blue feathers showed that these cavities have approximately spherical shape, with an average radius of 86 ± 13 nm and an average distance between centers of neighboring cavities of 240 ± 19 nm which quasi-periodical arrangement in the keratin matrix is responsible of the optical response observed in the plumage of male Swallow Tanagers (D’ Ambrosio *et al*., 2017; Skigin *et al*., 2019). To determine the mean radius (R) and the distance between neighboring holes (D) within the spongy matrix of the BL feather, we analyzed the microscopy images and obtained histograms for the aforementioned parameters (Figure 7). From these histograms it is found that R = 87 ± 13 nm, and D = 228 ± 31 nm. The mean R for both feather types is almost equal, while D is slightly smaller and more variable for the BL feather. The similarity between these values, and those of the BK feathers, is in agreement with the very similar reflectance spectra obtained with the micro-spectrophotometric measurements of both spongy matrices (Figure 4 (e)).

**Fig. 7.**
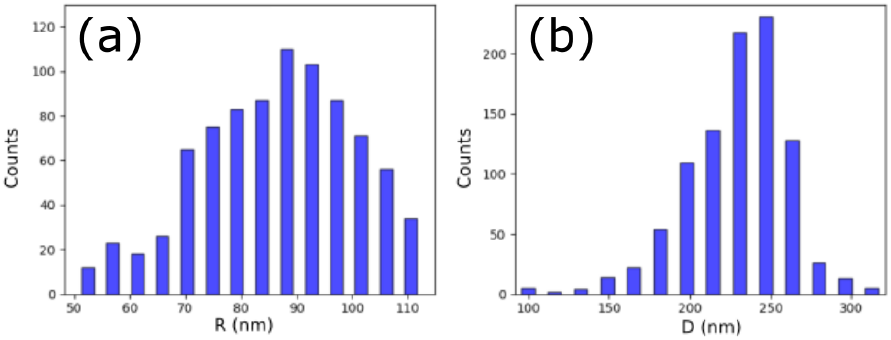
Histograms of (a) air cavities radius (R) and; (b) distance between adjacent cavities (D) within the spongy matrix of BL feather barbs.

## DISCUSSION

We compared the optical properties of the belly and back feathers of male Swallow Tanagers, which appear white and greenish-blue in the birds, respectively. We found that the belly feathers are not completely white when observed individually, instead they present a greenish hint at the tip of their most distal feather barbs. This was also captured by the reflectance spectra measured on the plumage, which showed an unsaturated reflectance peak at 550 nm, in agreement with the main reflectance peak of the back plumage. We performed a series of reflectance measurements to investigate the influence of the main components of the feather barbs in the optical response of the tissues. We corroborated the presence of a very similar spongy matrix in the tip of the feather barbs from both plumage patches. We also described the angle-dependence of the reflectance spectra, and the resulting color change, by measuring reflectance directly on the feathers with varying angle configurations of illumination-observation. Also, we found differences between feathers from both plumage patches in the shape and thickness of the cortex surrounding the feather barb, and in the deposition of melanin granules, both in feather barbs and barbules. Belly feathers showed a significant change in structure along feather barbs, and consequently in coloration, while back feathers were more uniform.

The reflectance spectra at the tip the exposed side of the barbs of both types of feathers here studied were quite similar, with both exhibiting a peak at approximately 550 nm. However, in the case of the back feather barb, the reflectance peak was more pronounced (i.e., more saturated). The reflectance for long wavelengths is low, approximately 20 % of the peak value. This intensification of the color purity could be produced by the presence of melanin granules in the inner side of the cortex, (i.e. below the spongy matrix), as suggested previously (Shawkey and Hill, 2006, Shawkey and D’Alba, 2017). Also, a well-defined reflectance peak is observed at the same spectral position (≈ 550 nm) when reflectance is measured directly on the spongy matrix of both feather types. This allows us to conclude that the hue of the plumage observed in a natural environment is mainly determined by the interaction of light with the spongy matrix, but melanin deposition and morphological variations alter the observed coloration considerably. Unfortunately, due to technical limitations we were unable to register reflectance in the UV range, which is visible to birds and where the greenish-blue plumage of male Swallow Tanagers has a secondary reflectance peak (Barreira *et al*., 2008; 2016). It would be interesting for future studies to compare the optical response of both types of feathers in the UV range. Given the similarities in the reflectance spectra produced by the spongy matrix of both feather types we would expect to also find this secondary reflectance peak on the belly feather barbs.

The reflectance of the back barbs differs substantially when they are illuminated from the exposed and inner sides. As observed in the microscopy images, the shape of the back feather barbs is triangular, therefore, it is not equivalent to illuminate the barb from one side or the other. In addition, the reflectance changes substantially between sides due to the presence of numerous melanin granules in the lateral and inner parts of the barb’s cortex and barbules. Given the high light absorption of melanin, the light intensity that reaches the spongy matrix in the case of inner side illumination is very low, and consequently so is its reflectance. In contrast, the reflectance curves for the belly barb corresponding to the exposed and inner side illumination setups are very similar. This suggests that the cortex thickness does not significantly influence the measured reflectance, given that the thickness of the belly feather barb cortex is very different between both, the exposed and inner sides. We found an engrossed upper cortex in these barbs that is likely involved in other functions other than plumage coloration (e.g., feather mechanical properties). We can also conclude that the absorption by keratin is negligible compared to that produced in the melanin granules, as stated before (Shawkey and Hill, 2006, Stavenga *et al*., 2011).

Highly white broadband reflectance (>70%) was described in the scales of white beetles of the genus *Cyphochilus*, generated by interconnected and randomly arranged chitin fibers with a diameter of about 250 nm (Vukusic *et al*., 2007). White coloration in birds plumage has been related directly to an anisotropic spongy matrix only for the crown feathers of *Lepidothrix isidorei* (Igic *et al*., 2016). A comparative analysis of white plumage across 61 species of birds of wide taxonomic coverage characterized how the macrostructures of the plumage (quantity, shape and density of barbs and barbules and internal structure of the barbs) affect the characteristics of the reflected light (Igic *et al*., 2018). The barbs of white feathers were in some cases thin and hollow, while others were thick and showed complex medullary layers inside. Species with complex internal medullary layers produced the higher reflectance values among white feathers of the studied species, but the specific contribution of the spongy keratin matrix was not determined (Igic *et al*., 2018). The keratin spongy matrix of feather barbs is thought to be produced through a passive, self-assembly, phase separation mechanism during feather growth (Dufresne *et al*., 2009). The size and inter-spacing of the air vacuoles in the spongy matrix have been shown to change among differently structurally-colored plumage patches (Stavenga *et al*., 2011, Tinbergen *et al*., 2013) indicating that birds can somehow regulate the size and periodicity of the spongy matrix during feather development to produce different color effects. However, a change from greenish-blue to white, as that of male Swallow Tanagers, would require an important increase in irregularity of the spongy matrix. We have not found such a change in the spongy matrix. Instead, we found that the belly plumage feathers had a similar spongy matrix at the tip of the feather barbs, without melanin granules, which gave the feathers a subtle greenish tint. The belly feather barbs near the rachis have large air voids and a flattened shape in accordance with previous observations (Igic *et al*., 2018), resulting in a flattened reflectance spectra. Overall, the macroscopic whitish effect of the belly plumage patch of male Swallow Tanagers seems to be the result of the over-stacking of feathers that lack of melanin underneath the spongy matrix and in their barbules, together with a change in the internal structure and barb shape closer to the rachis, and not a result of an increase of disorder in the spongy matrix itself.

The spongy keratin matrix was observed to be reduced in other white, gray and melanin-colored feathers of bird species known to produced quasi-periodic nanostructures in other structurally-colored plumage patches (D’Alba *et al*., 2012; Igic *et al*., 2016; 2018). This suggests that species that produce a quasi-periodic spongy matrix within their feather barbs with structural coloration, can also produce a similar structure in other plumage patches that do not show this type of coloration. The resulting plumage coloration may be therefore regulated by the juxtaposition of pigments and/or the reduction on the proportion of spongy matrix present in the barbs interior, rather than drastic changes in the periodicity and size of the internal nanostructures (Shawkey and Hill, 2006; D’Alba *et al*., 2012, Fan *et al*., 2019). Future studies, investigating color production of white or apparent pigment-based colored plumage patches on bird species with structurally-based coloration could help understanding if the presence of similar internal nanostructures in the feathers of differently-colored plumage patches is a common pattern. Also, female Swallow Tanagers show green and yellow plumage patches, which are thought to be the result of the combination of nanostructures and carotenoid pigments (Shawkey and Hill, 2005; Shawkey and D’Alba, 2017). It would be interesting for future studies to investigate the properties of the internal feather barb structures of female Swallow Tanagers in order to gain better knowledge about the regulatory processes leading to sexual dichromatism (Gazda *et al*., 2020) and the role of the elements that produce structural coloration in it.

## CONCLUSION

Overall, we have found that white and greenish-blue feathers of male Swallow Tanagers possess a very similar spongy matrix at the tips of their distal feather barbs, which also have almost equal reflectance spectra and show variation in the wavelength of peak reflectance with varying illumination-observation geometry. This feather internal nanostructure produces a subtle greenish coloration in the tips of the belly feathers. Even though this color effect is captured in the reflectance spectra of the plumage, it is very subtle and not visible (at least to humans) on the birds, perhaps as a result of high contrast with the surrounding greenish-blue colored plumage. The internal spongy matrix is reduced towards the rachis of the belly feathers barbs, flattening the reflectance curve and producing a real white reflectance spectra. Even though melanin contributes to the production of more saturated structural coloration, the sole presence of a quasi-periodical spongy matrix produces the differential reflectance of certain wavelengths defining the plumage color hue. The macroscopic white effect of the belly plumage patch is the result of the accumulated effect of the nanostructure variation across the feather barbs and the lack of melanin in the barbs and barbules. Our results highlight the importance of investigating the specific causes involved in changes in plumage coloration instead of making assumptions about the underlying mechanisms from the observed type of coloration only.

## Acknowledgements

We thank the technicians from the Sistema Nacional de Microscopia (MINCyT) that helped preparing the samples and obtaining the microscopy images. We also thank our colleagues from the División Ornitología of the Museo Argentino de Ciencias Naturales for their valuable comments on this study.

## Contribution

NCG, ASB, PLT, DS and MI designed the study, ASB selected the samples, TB and LM obtained the reflectance spectra and analysed the morphological data, DS, MI, TB, LM, NCG and ASB obtained the microscopy images, DS, MI and ASB wrote the manuscript, all authors reviewed the final version of the manuscript.

## Data availability

Data is available upon request to the authors

## Supplementary

Micro-spectrophotometry with varying angle configurations: To register the angular response of the feather we fixed the direction of the incident light and considered four feather orientations, as indicated in the central insets of Figure 8. Therefore, the angle of incidence is different in each configuration: *θ_i_* = 0°, 25°, 45° and 60°. Different observation directions were considered for each of these, as indicated in the diagrams with arrows of different shades.

**Fig. 8.**
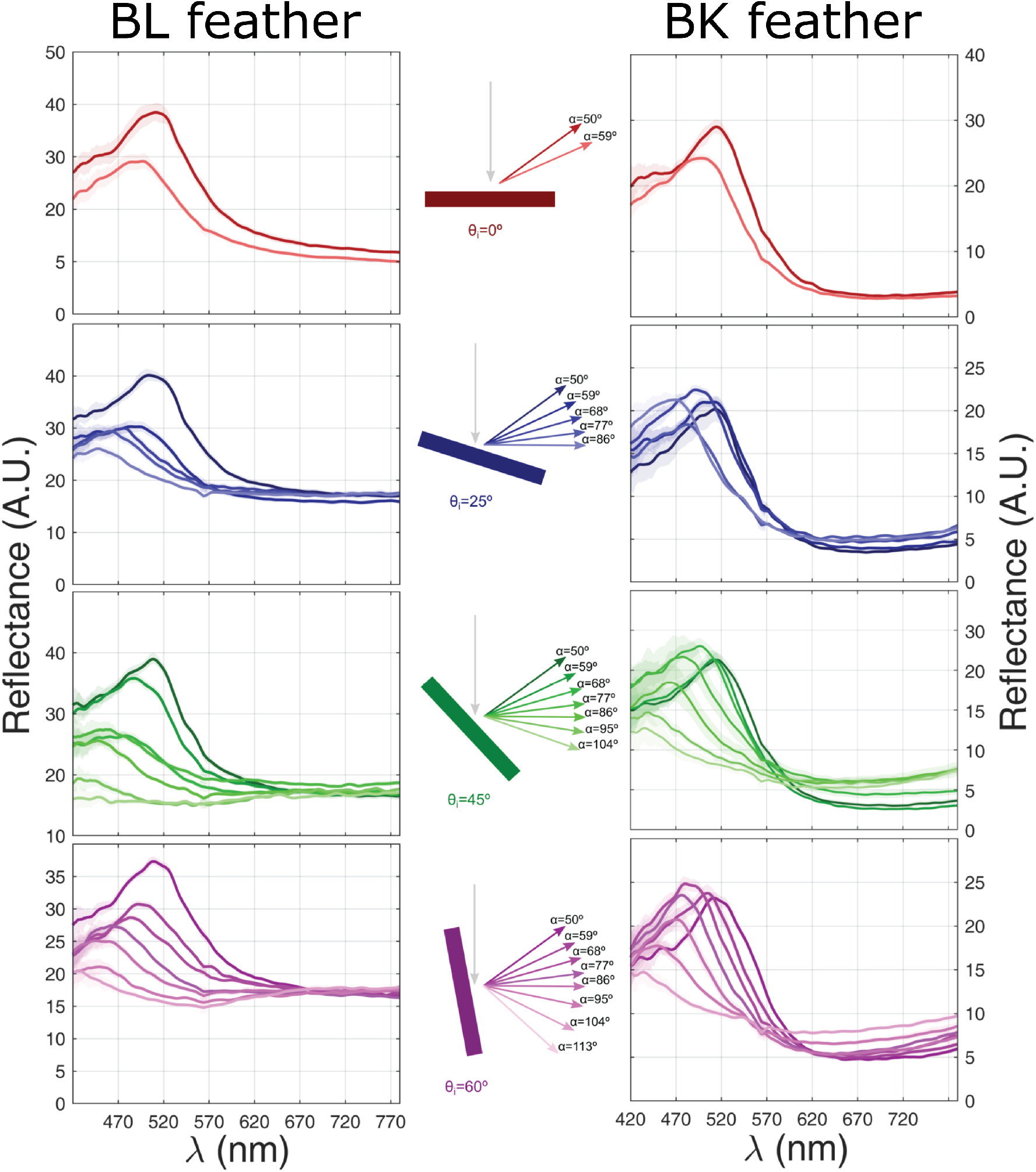
Angular reflectance measurements for the BL and BK feathers. Four angles of incidence were considered (*θ_i_* = 0°, 25°, 45° and 60°), as indicated in the central inset. The different curves in each panel correspond to mean reflectance (and standard error) for the different values of *α*, i.e, the angle between the incidence and observation directions.

